# playOmics: A multi-omics pipeline for interpretable predictions and biomarker discovery

**DOI:** 10.1101/2024.03.12.584088

**Authors:** Jagoda Głowacka-Walas, Kamil Sijko, Konrad Wojdan, Tomasz Gambin

## Abstract

**Background:** Multi-omics analysis is increasingly popular in biomedical research. While promising, these analyses confront challenges in data integration, management, and interpretation due to their complexity, diversity, and volume. Moreover, achieving transparency, reproducibility, and repeatability in multi-omics analyses is essential for facilitating scientific collaboration and validation of complex datasets.

**Results:** We introduce playOmics, an open-source R package tailored for omics data analysis. It facilitates data management and biomarker discovery through various visualizations, statistics and explanations for boosted interpretability. playOmics identifies significant prognostic markers and iteratively constructs logistic regression models, identifying combinations with high predictive performance. Our tool enables users to make direct, model-driven predictions by inputting new data into the selected pre-trained model. playOmics performed well in handling extensive datasets and missing data, showing a mean validation MCC of 0.773.

**Conclusions:** playOmics demonstrates the balance between model complexity and interpretability, crucial in biomedical research for understanding model decisions. playOmics’ approach promotes a flexible model selection process, encouraging exploration and hypothesis generation in biomarker discovery. The dockerized setup and intuitive graphical interface of playOmics support its adoption in a wide range of research and clinical settings, adhering to principles of open science, enhancing reproducibility and transparency.

## Introduction

### Background

Multi-omics studies, encompassing diverse layers of biological information such as genomics, transcriptomics, proteomics, and metabolomics, are pivotal in dissecting complex molecular signatures associated with specific phenological traits or disease states [1]. However, despite its potential, the true value of comprehensive omics analysis often encounters significant challenges, particularly in data integration, management and interpretation [2, 3]. These challenges arise due to the complexity, diversity and voluminous nature of the data, posing substantial obstacles in effectively utilizing integrated omics analysis.

One of the primary challenges in multi-omics data analysis is the scenario where the number of features (p) greatly exceeds the number of observations (n). This imbalance requires robust techniques such as dimensionality reduction or feature selection to ensure that results are meaningful and interpretable [4]. This issue is further extended in rare disease studies, where patient samples are deficient, emphasizing the need for methods capable of extracting maximum information without overfitting [5].

Effective data management, encompassing preprocessing and cleaning, is essential for harmonizing data from various sources. This process ensures data quality and compatibility, which is crucial for downstream analysis. Various approaches to data integration have been proposed, with each having its own advantages and disadvantages [6, 7]. The most common approach, early integration (also known as data concatenation), involves combining different datasets or omics layers before any analysis, allowing for simultaneous analysis of all data types. Despite its ease of application, early integration risks overlooking the complex relationships between different omics layers inherent in biological systems, as it assumes a straightforward additive effect of combining datasets. Hierarchical integration, on the other hand, enriches the analysis by incorporating regulatory relationships between omics layers, informed by prior knowledge and external sources like databases and literature. However, by focusing on known relationships, novel interactions might be overlooked as they are not yet represented in databases or literature.

### Transparency, reproducibility and repeatability in multiomics analyses

The complexity and scale of comprehensive omics data demand research methodologies and findings to be transparent, reproducible, and repeatable. Ensuring such consistency not only builds trust in the findings, but also facilitates scientific collaboration, allowing other researchers to validate and build upon existing work [8, 9]. Achieving reliability is fundamental to validating the results and conclusions drawn from complex datasets [10]. While the quintessential principles of open science — transparency, accessibility, and reproducibility—are often addressed through clear methodological guidance offered by various analytical packages, the tools known from machine learning engineering, which includes streamlined operation on modeling pipelines, model versioning and monitoring, is not always integrated in the traditional tools. Additionally, the capability to adjust models across various datasets marks the robustness and adaptability of analytical tools, significantly increasing the practical application of research outcomes.

Another important aspect is the management and validation of results. This requires robust statistical methods and practices to confirm the reliability of the findings. Permutation experiments, for instance, are a well known technique used to determine the significance of the results [11]. The adoption of metrics specifically aligned with the nature of the data, including those for addressing issues like imbalanced datasets, further confirms the findings’ validity [12].

### Interpretability

Interpretability, a critical factor in fields with significant decisionmaking consequences like healthcare and finance, has become crucial in the field of multi-omics data analysis. As multi-omics experiments grow in complexity, there is an increasing demand for models that are not only accurate, but also transparent in their decision-making processes. Techniques such as feature importance analysis, partial dependence plots and SHAP (SHapley Additive ex-Planations) values are instrumental in elucidating how individual features influence predictions on a local scale, providing insights into specific data points or predictions. In contrast, global explanators, including interpretable models like decision trees and linear models, allow for an understanding of the model’s behavior as a whole [13, 14]. Integrating interpretability into model development in multi-omics not only boosts trust and transparency, but also ensures that models can be effectively and responsibly applied in critical areas like clinical decision-making, thereby bridging the gap between complex data analysis and its real-world healthcare applications.

### Current approaches

The multi-omics data analysis field has evolved through the development of tools designed to tackle specific aspects of data integration and analysis. This progress has been driven by efforts to create machine learning methods that automatically integrate omics data. To name some, broadly applied R packages like mixOmics, MOFA, and iCluster have played a significant role in this development. mixOmics, a universal package for both supervised and unsupervised analysis due to its implementation of methods like LDA, PCA, and CCA, and the DIABLO framework, is widely used for cancer subtype characterization and disease association studies [15, 16]. MOFA employs factor analysis for dimensionality reduction, aiming to identify common factors across various omics layers that account for the greatest data variance; its effectiveness has been proven in survival analysis and drug response prediction [17]. The iCluster utilizes joint latent variable models to reduce dimensionality and integrate data across omics layers, and have been effectively applied in identifying cancer subtypes and patient stratification [18]. More recently, the Python library QLattice has been introduced, employing a symbolic regression approach to generate simple, predictive models from omics data, where complexity is moderated by a Bayesian criterion [19]. It’s particularly suited for clinical decision-making and patient care. However, despite its direct interpretability, access to QLattice is governed by specific licensing terms, and its source code is not openly available, which may limit its adoption in research environments that prioritize open science principles and collaborative development. To the best of our knowledge, no tool combines the ease of analysis tracking with the provision of user-friendly graphical interface and interpretable models serving.

Addressing existing gaps, we developed playOmics, an R package specifically designed for multi-omics analysis. Our strategy with playOmics was to simplify the process of integrating complex omics data, facilitating the identification of prognostic markers and development of effective predictive models (see 1 for a comparison with existing tools). We prioritized ease of the data processing and model engineering, introducing an interactive GUI for model exploration, serving and validation. playOmics emphasizes model interpretability through various statistics, visualizations and the use of local explainers like SHAP values for in-depth analysis.

## Implementation

### Overview

The playOmics package was developed using R Statistical Software [20] and builds on a number of existing packages, in particular, tidyverse [21], tidymodels [22], mlr3 [23], shiny [24] and DALEX[25]. Its source code and documentation are accessible on GitHub, and can be found at [26]. To facilitate the usage and reproducibility, we have prepared two distinct docker images: one encompassing a complete test environment with RStudio, and another containing test data. A docker-compose file, along with a script for testing purposes, has been curated and hosted on GitHub [27]. This arrangement provides an isolated, self-contained setup for running the playOmics environment.

A graphical abstract of the data pipeline proposed in the playOmics package is presented in Figure 1. In the following sections, the main steps of the analysis will be discussed in details.

**Figure 1.**
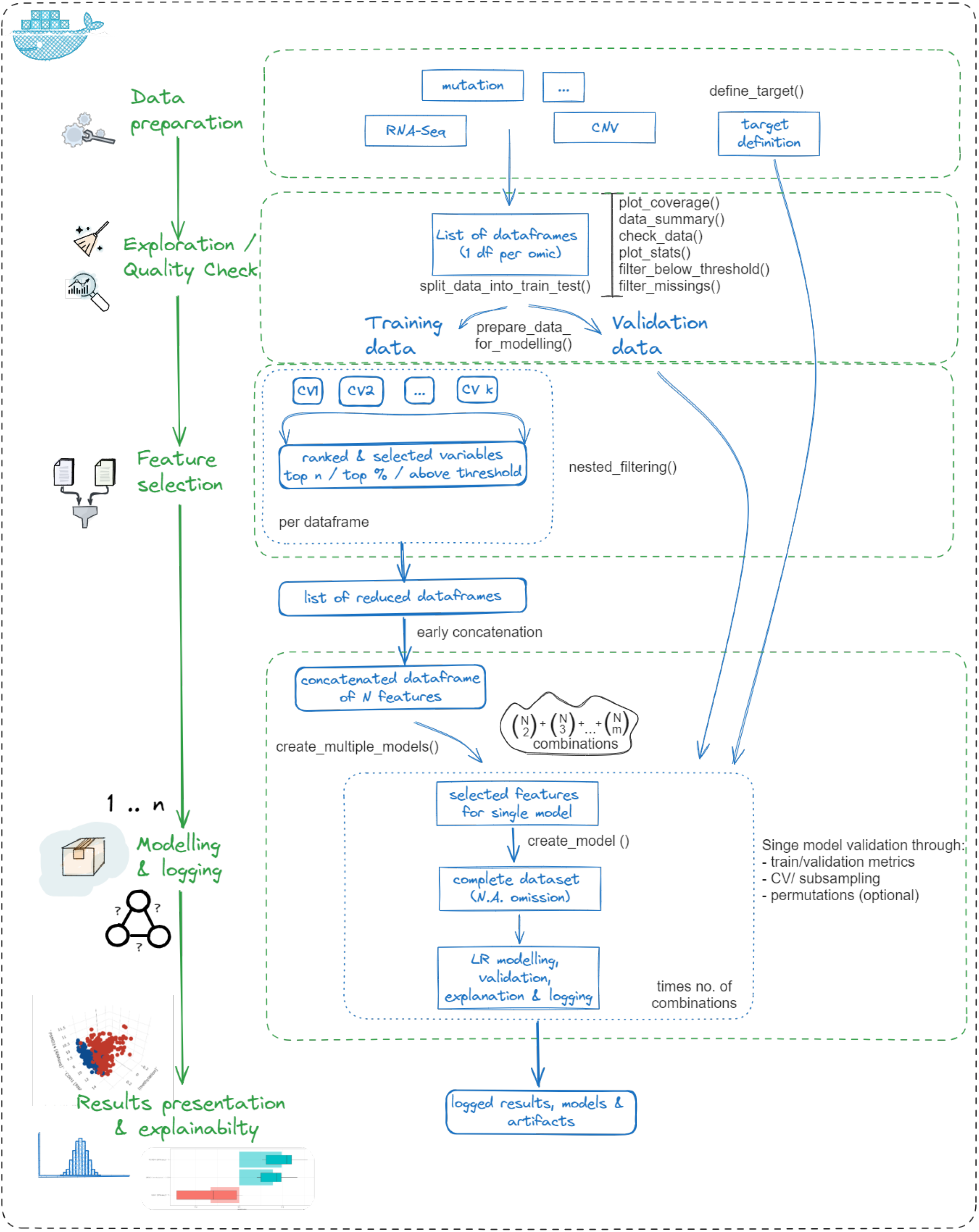
Schematic overview of playOmics, presenting the main steps of the analysis (left side, green font) together with their corresponding data products and associated functions (right side, blue and black font, accordingly). The initial steps involve data preparation, exploration and quality check, ensuring that datasets adhere to a common structure to simplify manipulation and preprocessing. In the next step, feature selection in a cross-validated manner is performed separately for each dataframe, which allows for a balanced contribution to the modeling process. The subsequent step is central to playOmics, involving the training, evaluation and logging of models. The final step offers a module for result presentation and interpretation. Within playOmics, a graphical user interface helps in model management, while a range of statistics and visualizations assist user in the results interpretation. The workflow is available as a standalone package or as a dockerized environment.

### Data preparation

In the initial phase of our omics data analysis, we focus on integrating various data types. These should be formatted as data frames suitable for downstream analysis and follow tidy data principles, with observations in rows and variables in columns. Here, the variables correspond to measurements from different omics types in continuous or binary scales. This format promotes consistency across all data types, crucial for further unification and analysis. Our approach involves storing the different data in a list format, which simplifies the manipulation of individual datasets and supports a unified view of the aggregated data.

The definition of the analysis target guides the way how we treat the data in the preprocessing step. We use phenotype data to set the prediction objective (e.g. patient survival status) and to point out common observation identifiers, essential for merging data across different datasets. This stage also includes the possibility of selection of the “positive” class - the outcome we aim to predict. After a target is set, we proceed to convert all remaining factor and character variables into a binary format using one-hot encoding. Additionally, each variable is tagged with an identifier that links it back to its original dataframe, ensuring traceability throughout the analysis process.

### Data exploration and quality check

In the exploration stage, we assess the alignment of different datasets. We begin by conducting a visual examination of the extent of overlap across different data types using upset plots to assess the shared data points among our datasets. This process helps us identify any datasets with insufficient overlap, allowing us to consider the removal of any modality that lacks adequate coverage. Next, we conduct a detailed quality check of all the data. This includes examining basic statistics for both numerical and non-numerical data to uncover necessary adjustments, such as converting text data to numeric formats and spotting columns with many missing values or showing little to no variation. Identifying these issues allows us to adjust data for consistency across the datasets or to eliminate variables that, due to excessive missing values or lack of diversity, may not contribute valuable insights to further analysis.

The playOmics package allows for the application of omic-specific standards in data preprocessing, which is crucial for managing the diversity and complexity of omics datasets. This can refer to a variety of methods, which is beyond the scope of the package.

However, addressing the common challenge of data filtering, we have implemented two general functionalities: one for filtering out values below threshold and another for removing variables with a high percentage of missing values, following the thresholds for acceptable levels of missing or low-quality data set by user.

The segregation of data into training and test subsets is another step toward the modelling process. This split is conducted on each dataframe separately. Stratified sampling on a phenotype dataframe is performed and, subsequently, the dataframes corresponding to different omics are split based on the same subsets of train and test IDs.

### Feature selection

Feature selection holds a pivotal role in the omics data analysis due to the high-dimensionality typically associated with such datasets. Our approach is to conduct feature selection on the training data separately for each dataset, ensuring balanced contribution to the modeling process and preventing any single dataset from dominating.

We adopt a method that involves nested filtering, where features are ranked and selected based on their relevance to the analysis in the univariate manner. This is achieved by running cross-validated feature ranking, which reduces the risk of overfitting in the selection process and then averaging the ranks to determine the most relevant features. Ranking is performed using the mlr3filters package, which offers a range of evaluation metrics [28].

Based on the mean results of cross-valitation runs, we then select the set of best features for modeling. The feature selection methods include: ‘top n’ for selecting a defined number of features from each dataset (a default), ‘percentage’ for choosing a certain proportion of variables from the dataframe, and ‘threshold’ for picking features that exceed a specific mean score.

In our approach, we follow an early concatenation strategy. After feature selection step, we combine all preprocessed datasets into a single, unified dataframe which is further utilized in models’ development.

### Modelling

In our modeling approach, we construct a range of logistic regression models tailored for supervised binary classification, each using different combinations of predictors. Our goal is to explore all possible combinations to identify the most effective set of features that distinguish between the two groups. To achieve this, we systematically create a set of feature combinations that need to be evaluated. The number of possible unordered combinations of a specific size (m) that can be selected without replacement from a larger set of available features (N) is determined as follow:

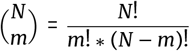

Where: N is the total number of available features and m is the number of predictors (independent variables) selected for a specific combination.

Logistic regression strikes a balance between complexity and explainability, making it particularly suitable for applications where interpretability is crucial. This method models the probability of an event, such as the clinical outcome. If the probability is higher than the defined threshold, then the model classifies the outcome as “positive” class. For our logistic regression models, we carefully limit the number of features, especially in small sample sizes, to prevent overfitting. A logistic regression binary classification algorithm to generate a m-variable predictor is defined as following:

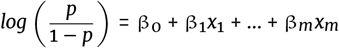

Where: p – expected probability, *x*_1,..,*m*_ – independent variables, β_0,…,*m*_ – regression coefficients.

One of the critical aspects of forming these models is the careful handling of missing data. Rather than removing missing data early on, we delay this step until the construction of individual models. This approach is beneficial in omics data analysis, given the complexity of aligning various omics layers that range from hundreds to tens of thousands of variables. Our approach ensures that we maximize the use of available data and avoid discarding potentially significant signals, especially in the contexts of rare disease studies, where each data point can be critical.

### Results presentation and interpretability

In playOmics, we prioritize interpretability to enhance the understanding of complex multi-omics data. Therefore we developed a graphical interface to facilitate the management and interpretation of experiment results. It includes various analytical panels for a deeper examination of the experiment, including the analysis summary, detailed views of variables’ statistics and visual explanations of single model predictions 2. Primarily, the Main Panel displays a scored list of predictive models from the analysis, offering key identification information and their metrics for the training and validation sets. It also facilitates quick navigation to other sections through model-specific buttons (Fig. 2A). The Analytes Overview Panel provides the general statistics for individual molecules, along with their average metrics across all models to which they contributed, such as MCC, F1, ROC AUC, NPV/PPV and more, offering insights into the quality of molecular contributors within the experiments (Fig. 2B). The Experiment Overview Panel enables the visualization of selected experimental metrics through histogram plots, offering an overview of the outcomes (Fig. 2C). The Single Model Overview Panel displays the training data and its visual representations on 2- and 3-dimensional plots, supplemented with statistical information, i.e. median and IQR values within each group, which helps to identify patterns and understand the predictors’ roles in distinguishing the target status (Fig. 2D).

**Figure 2.**
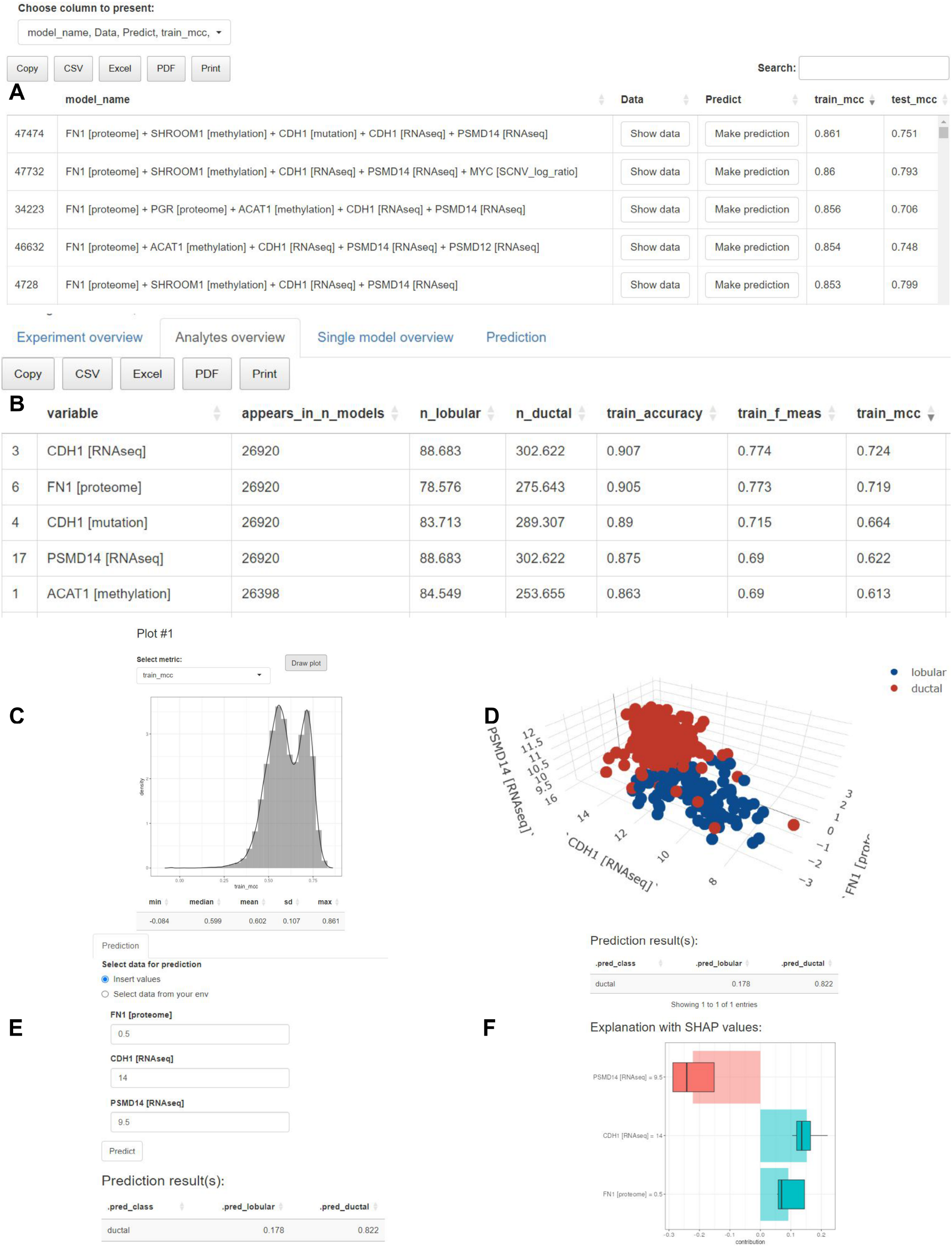
User interface components in playOmics. A. Result panel presents a list of predictive models. Selected performance metrics are displayed alongside each model for evaluation. Two primary actions can be initiated from here: ‘Show data’ button leads to a detailed view of the model’s input data, and ‘Predict’ button opens the prediction panel for the selected model. B. The Analytes Overview Panel displays statistics for each molecule and their average metrics across contributing models C. The histogram of training MCC values among all scored models with the corresponding statistics. D. A three-dimensional scatter plot representing a best-performing 3-variable model consisting of CDH1, PSMD14 (RNA-Seq data), and FN1 (proteomic data), with a training MCC of 0.815 and validation MCC of 0.734. Each data point represents an observation, while color distinguishes between different outcomes (blue dots - lobular subtype, red dots - ductal). E, F. In the prediction panel, users can input new data for the model of interest either by manually entering values or by selecting an existing dataframe from the environment. Upon submitting the input data for prediction, the interface returns a table that includes the probabilities of each class along with the final predicted class. To facilitate explainability, SHAP (SHapley Additive exPlanations) values are provided as visualization, representing the contribution of each feature to the prediction outcome. The blue color corresponds to lobular class, while red corresponds to ductal class.

Additionally, the Prediction Panel enables direct predictions with the selected model. After the initial phase of model training and logging is completed, users can input new data into the chosen model to receive immediate, model-driven predictions (Fig. 2E).

An important component of our interpretability approach is the use of SHAP values in the Prediction Panel (Fig. 2F). SHAP values are a powerful tool for local model explanations, revealing how individual features in a model affects specific predictions. They help to quantify the impact of each feature on the model’s output, pointing out why the model makes certain decisions for each specific case. This is particularly valuable in clinical scenarios, where understanding the factors driving a model’s predictions can be crucial for decision-making process and elevate trust.

### Validation

playOmics incorporates permutation experiments as a validation method to ensure the robustness of our models, accessible through Single Model Overview Panel. These experiments assess the relationship between the target variable and the predictors. By repeatedly retraining the model on these permuted datasets (i.e. permutation of classes) and evaluating their performance, we construct a null distribution that reflects what one might expect under the assumption of random labeling. The observed performance of the selected model is then rigorously compared against this null distribution. A known limitation of permutation experiments is the practical challenge of exploring all possible combinations, especially as the total number of observations increases. This is represented mathematically as 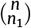, where n is the total number of observations and *n*_1_ represents the number of observations in one group, with the total being the sum of *n*_1_ and *n*_2_. Therefore, permutation tests are particularly useful for studies with a smaller sample size. Marozzi et al. [29], based on comprehensive literature reviews and simulation studies, suggests a practical approach for permutation tests, recommending 5000 for actual applications where the significance level is 5% or 10000 for the significance level 1% to achieve reliable p-value estimates, whereas Legendre&Legendre [30] suggest 500-1000 permutations during an exploratory data analysis and 10000 for final results.

### Reproducibility

playOmics enhances the reproducibility of experiments through extensive logging capabilities that encompass both overarching experiments and specific models. Every aspect of the machine learning process, from parameters to results, is recorded in userdefined directories. This structured approach also applies to the management of artifacts. Input datasets, models and explainers are stored in binary formats facilitating their reuse on new data sets or in-depth analysis for greater insight. Furthermore, analysis parameters, model configurations, logs, and performance metrics are stored as JSON files to support efficient troubleshooting and enable precise replication of experiments. Its Docker-based setup allows smooth in various environments and promotes collaboration among researchers.

### Performance evaluation

In our analysis process, the computational efficiency and resource requirements are critical considerations, especially when dealing with a growing number of variables. The performance is primarily driven by two factors: the maximum number of variables in a single model (m) and the number of variables selected for the modelling experiment (N).

As the number of variables per model increases, so does the computational load (Fig. 3A). This is due to the exponential increase in the number of combinations that need to be evaluated when more variables are involved. Models with multiple predictors require an extensive exploration of all possible combinations, starting from simpler ones and progressing to more complex arrangements.

**Figure 3.**
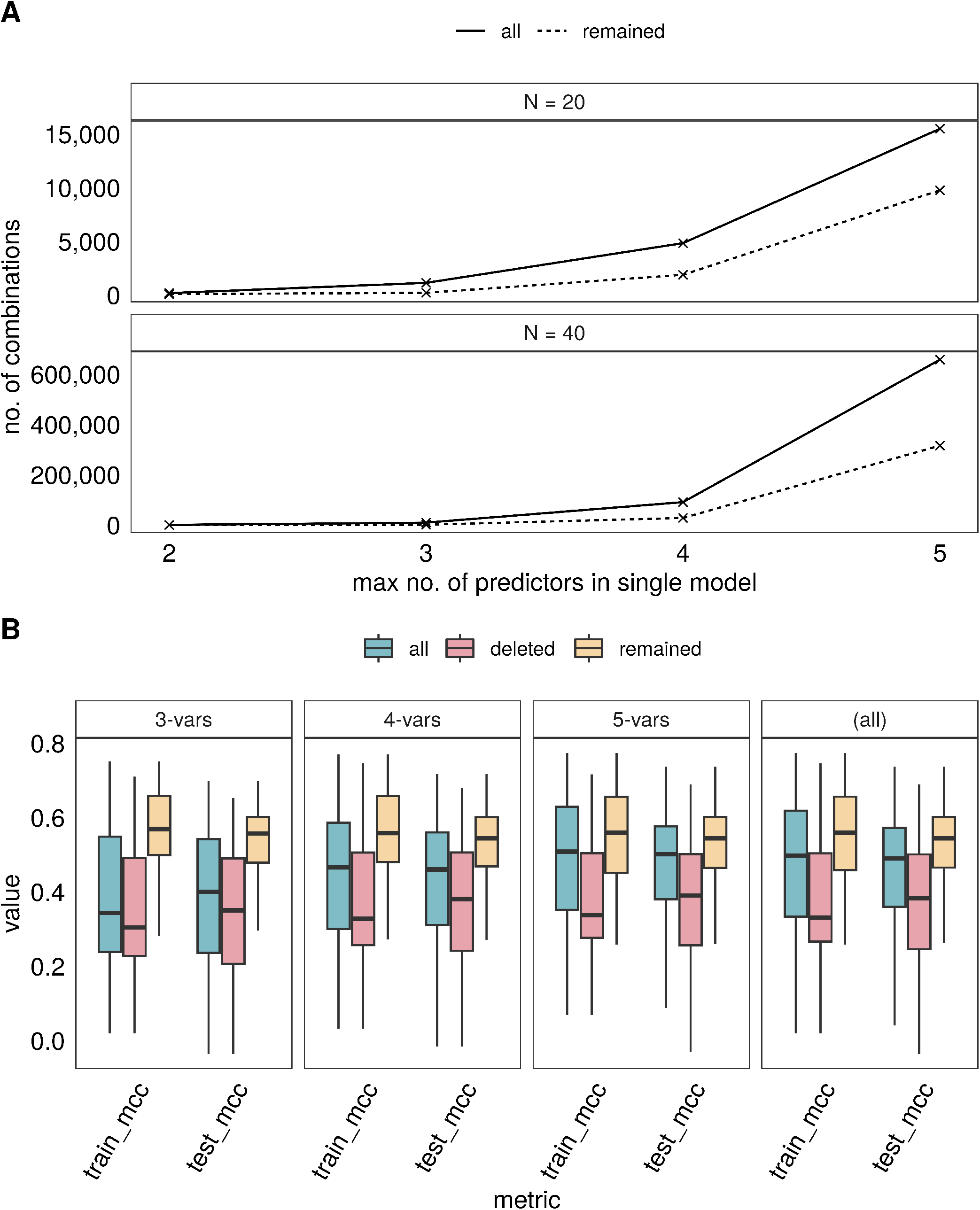
Impact of model removal on the number of final models and MCC metrics. A. The plot displays the count of final models resulting from varying the maximum number of predictors allowed in a single model and the numbers of available predictors in the final set (N), demonstrating the effect of model pruning when the threshold for training MCC is set below 0.3. Given that MCC values range from -1 to 1, where 0 indicates no better than random chance, setting the threshold at 0.3 ensures that only models with a certain level of meaningful predictive ability are retained. Solid lines represent the total number of models before elimination, while dashed lines indicate the remaining models after trimming. B. The distribution of MCC values for experiments with 20 features (N = 20), categorized by the post-removal models’ status. Colors denote the full set of models (“all”), models that were retained after pruning (“remained”), and those that were excluded (“deleted”). The boxplots illustrate the median MCC (center line), interquartile range (box edges), and the range within 1.5 times the interquartile range from the upper and lower quartiles whiskers).

Furthermore, the total number of variables chosen for the experiment plays a significant role in computational intensity. Each additional variable escalates the complexity of combination generation, demanding more computational power. Therefore, careful selection of variables and appropriate setting of variable limits are essential. By strategically managing these aspects, we can effectively balance computational demands, ensuring the efficiency of the process without compromising the depth and accuracy of our analysis.

To enhance the efficiency of our analysis, playOmics is able to remove less effective models. This means we actively delete models that don’t meet standards of performance defined by the user, therefore the same combination of predictors will not form models of higher order. Such a practice allows us to focus on models that demonstrate the highest predictive power. It works as follow: initially, models built with pairs of features (two-element models) are assessed. If these models show weak predictive performance, based on pre-established metrics and threshold, they are excluded from further analysis. The rationale behind this strategy is that if two variables do not significantly contribute to predicting the target, their inclusion in a three-element model is also likely to be ineffective. This continues until the defined maximum number of features allowed in the model.

This approach has been experimentally validated, proving effective in dealing with the complexities of large datasets and ensuring that our analysis is driven by the most robust and informative models (Fig. 3B).

### playOmics evaluation

To demonstrate the capabilities of the playOmics package and conduct comparison with other tools, we utilized the Breast Cancer (BRCA) dataset from The Cancer Genome Atlas (TCGA) project. Our analysis focused on predicting histological subtypes: infiltrating ductal and lobular carcinoma. However, although coming from the same source, it is important to note that the data vary between two sections of the results chapter.

In the use case section, we employed what we refer to as the “extensive” dataset, downloaded via the LinkedOmics portal [31]. The annotation data were available for 1097 subjects. We incorporated the following datasets in our study: clinical data (20 features), proteome (176 features), methylation (20107 features), miRNA (824 features), mutation (7967 features), RNA-Seq (20156 features), and CNV (24777 features).

To benchmark our work against existing analytical methods, we conducted a comparative analysis assessing the performance of the playOmics algorithm alongside several established alternatives. This analysis includes comparisons with the QLattice package, the autoML framework, Lasso regression, and decision tree algorithms. Lasso regression, decision trees, and autoML were selected as state-of-the-art algorithms, with Decision Trees serving as a baseline for interpretable models. The QLattice tool was selected due to its similarities in model design project - it utilizes relatively small models, which makes it comparable to playOmics goal. However, comparisons with other algorithms listed in Table 1, such as mixOmics, MOFA, and iClusterPlus, were not feasible due to significant methodological differences, such as a lack of ability to perform supervised classification.

**Table 1.**
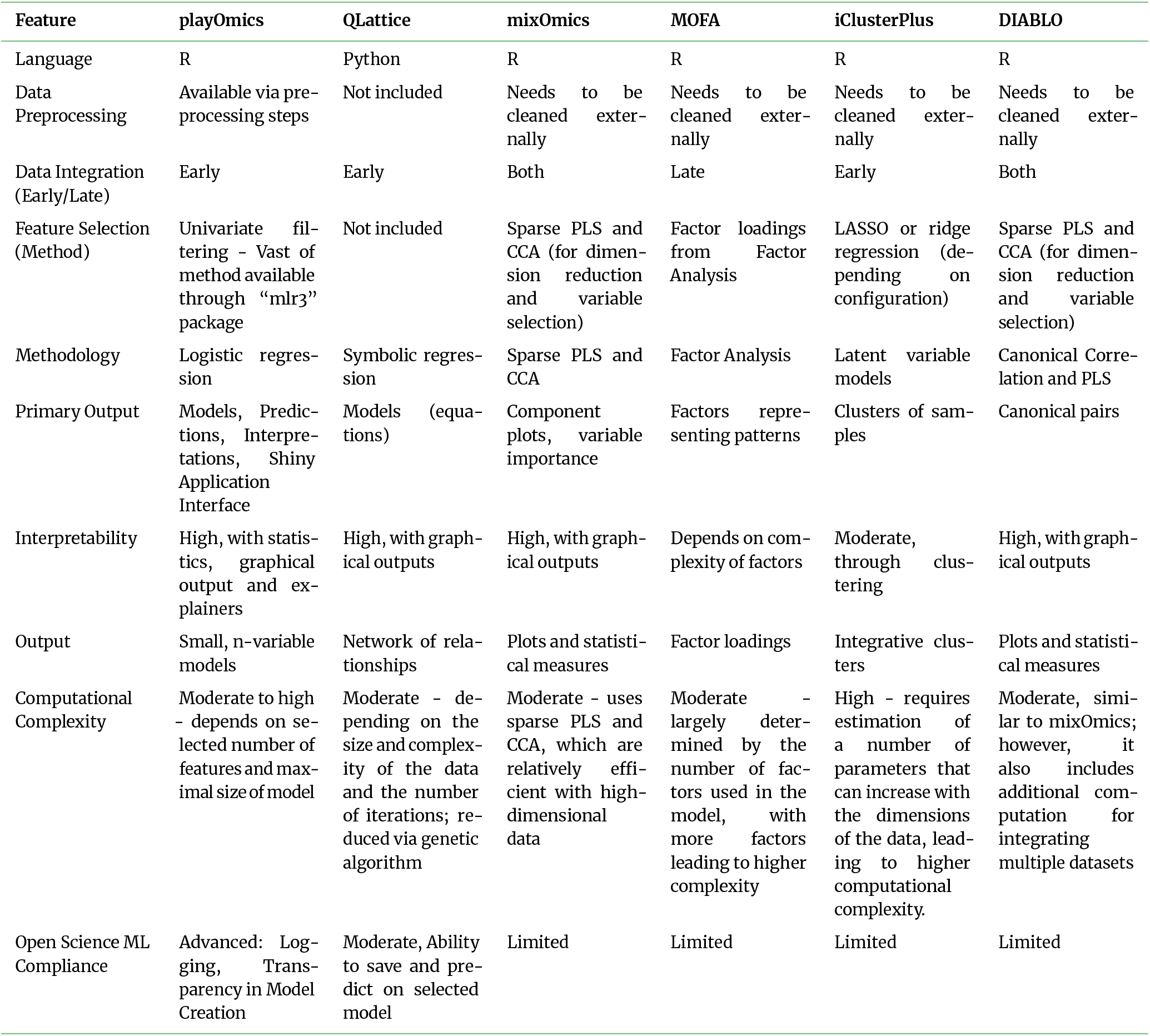
Comparative Overview of Multi-Omics Analysis Tools.

| Feature | playOmics | QLattice | mixOmics | MOFA | iClusterPlus | DIABLO |
| --- | --- | --- | --- | --- | --- | --- |
| Language | R | Python | R | R | R | R |
| Data Preprocessing | Available via pre-processing steps | Not included | Needs to be cleaned externally | Needs to be cleaned externally | Needs to be cleaned externally | Needs to be cleaned externally |
| Data Integration (Early/Late) | Early | Early | Both | Late | Early | Both |
| Feature Selection (Method) | Univariate filtering - Vast of method available through “mlr3” package | Not included | Sparse PLS and CCA (for dimension reduction and variable selection) | Factor loadings from Factor Analysis | LASSO or ridge regression (depending on configuration) | Sparse PLS and CCA (for dimension reduction and variable selection) |
| Methodology | Logistic regression | Symbolic regression | Sparse PLS and CCA | Factor Analysis | Latent variable models | Canonical Correlation and PLS |
| Primary Output | Models, Predictions, Interpretations, Shiny Application Interface | Models (equations) | Component plots, variable importance | Factors representing patterns | Clusters of samples | Canonical pairs |
| Interpretability | High, with statistics, graphical output and explainers | High, with graphical outputs | High, with graphical outputs | Depends on complexity of factors | Moderate, through clustering | High, with graphical outputs |
| Output | Small, n-variable models | Network of relationships | Plots and statistical measures | Factor loadings | Integrative clusters | Plots and statistical measures |
| Computational Complexity | Moderate to high - depends on selected number of features and maximal size of model | Moderate - depending on the size and complexity of the data and the number of iterations; reduced via genetic algorithm | Moderate - uses sparse PLS and CCA, which are relatively efficient with high-dimensional data | Moderate - largely determined by the number of factors used in the model, with more factors leading to higher complexity | High - requires estimation of a number of parameters that can increase with the dimensions of the data, leading to higher computational complexity. | Moderate, similar to mixOmics; however, it also includes additional computation for integrating multiple datasets |
| Open Science ML Compliance | Advanced: Logging, Transparency in Model Creation | Moderate, Ability to save and predict on selected model | Limited | Limited | Limited | Limited |

To enable the comparative analysis, we adopted a second dataset, which we will refer to as a ‘limited’ dataset, distinct from the one used in the use case section. This dataset, originally employed in the QLattice publication and accessible through Github [32], was selected for its absence of missing values. It consists of four specific datasets (CNV - 860 features, proteome - 223 features, RNA-Seq - 604 features, mutation - 249 features) and covers 705 patients. This dataset’s lack of missing values is key for our benchmarking, especially for comparison with QLattice and Lasso regression algorithms, which cannot process missing data. Choosing it avoids the problem of missing data being excluded when datasets are combined, which would otherwise lower the number of observations and affect the fairness of our comparison. A sample weight approach was adopted on this data due to higher class imbalance in comparison to the extensive dataset (about 4.5:1 in the proportion of ductal to lobular in the limited dataset, compared to about 3:1 in the extensive dataset).

## Results

### Use case

To illustrate the typical workflow within playOmics, we conducted a detailed step-by-step analysis using an extensive dataset as an input. The complete script for this analysis is available on Github, and the results are presented in Figure 2.

The clinical dataset initially included nearly a thousand patients, however, there was significant variance in data coverage across different datasets and variables. At most, information was available for 379 patients across all datasets. Specifically, the minority class (lobular cancer) ranged from 50 to 145 samples in the training set and from 13 to 37 in the validation set. This discrepancy in data availability added an additional layer of complexity to our analysis, challenging for many algorithms that do not accommodate missing data.

In the filtration phase, a univariate variable selection was conducted using the Area-under-the-Curve (AUC) metric to evaluate the distinction between lobular and ductal samples. We selected top 5 features from each dataset, except for clinical data, which resulted in a total of 30 features for modelling experiment. Subsequently, these analytes were integrated into logistic regression models with a maximum of 5 variables. In total, 157,387 models were scored, and 16,607 were eliminated in the early reduction phase. The computational time for the experiment reached 4.5 hours, with parallel processing across 10 cores and approximately 10 GB of RAM.

For the overall experiment, the average training MCC stood at 0.602 (with a standard deviation of 0.107), whereas for the validation set it was slightly lower at 0.589, (with a standard deviation of 0.152) (Fig. 2C).The highest-performing model included five variables: CDH1 and PSMD14 from the RNA-Seq dataset, FN1 from proteome dataset, SHROOM1 from methylation data, and CDH1 from mutation data, achieving an MCC of 0.861 on the training set and 0.751 on the validation set.

Another approach to interpreting the data is to analyze molecules and their respective statistics across all evaluated models, available through the Analytes Overview Panel (Fig. 2B). For instance, CDH1, highlighted previously as a component of the best model, stands out as a key molecule, with its RNA-Seq expression levels averaging a training MCC of 0.724, highlighting its importance in distinguishing between lobular and ductal cancer subtypes. Its mutation data further supports its significance, registering an MCC of 0.664. The presence of CDH1 mutation, as well as its lower expression, corresponds to lobular subtype, what was primarily found in the original publication describing BRCA dataset [33] and confirmed in our study (Fig. 2D).

To further emphasize interpretability of playOmics, in Figure 2D we demonstrate the outcomes of a best-performing 3-variable model that incorporates FN1 from the proteome dataset and two RNA-Seq markers, CDH1 and PSMD14. While this model exhibits slightly reduced performance metrics—0.815 in training and 0.734 in validation MCC—compared to a 5-variable model, it outperforms this more complex model in terms of simplicity and clarity. The 3D visualization captured in Figure 2D helps to identify the differences between cancer subtypes, illustrating how the combined influence of these three variables builds a robust predictive model.

Furthermore, we conducted direct predictions using this model to demonstrate its practical application, with the results depicted in Figures 2E and 2F. When presented with new data, the model estimated a 17.8% probability for classifying the sample as lobular and an 82.2% probability for ductal classification, resulting in the assignment to ductal class. The SHAP values calculated for this prediction reveal the individual contributions of each variable: CDH1 and FN1 shift towards a ductal classification, whereas PSMD14 leans the prediction towards lobular. This nuanced understanding of variable impact underscores the model’s interpretability, aiding users in making informed decisions.

### Comparison with other tools

The results presented in 2 provide a comparison of the performance of the playOmics algorithm against other established algorithms for BRCA subtypes (ductal/lobular) prediction on a limited dataset. The comparison utilized the Matthews Correlation Coefficient (MCC) metric to evaluate model performance on the validation dataset, with additional insights provided by training data metrics (indicated within brackets).

playOmics demonstrates a reasonable validation performance, especially when features were selected using MIM (0.683) and a bit lower for the other two methods (AUC 0.663 and MRMR 0.634). QLattice, on the other hand, consistently shows high MCC values across all selection methods, peaking with MRMR (0.715). The autoML model with the top 5 features from all data shows the highest validation MCC (0.764 with AUC, 0.771 with MRMR) among all models, indicating that it can make highly accurate predictions. Lasso’s performance is generally strong but does not reach the peak performance of autoML. Decision trees, selected as a baseline for interpretable models, show lower performance compared to other models, suggesting that this approach might be less effective for this particular prediction task.

When comparing models built from a larger set of features, specifically 40 features, against those constructed from a smaller subset, the impact on predictive performance rises for the number of features decreasing. For example, autoML models utilizing the top 10 features from each dataset (40 features in total) show varying performance with validation MCC scores ranging from 0.666 to 0.652 when selected by AUC and MIM, respectively, while the model with the top 5 features from all data achieves the highest validation MCC scores of 0.764 and 0.771 when features are selected by AUC and MRMR, respectively.

## Discussion

In this work, we introduce playOmics, a modeling pipeline developed to improve interpretability in omics data analysis. The first version of this pipeline have been applied in previous work, allowing to identify biomarker combinations that distinguish children with epilepsy status in a small population affected by TSC mutation [34].

Our aim with playOmics was to simplify the management of diverse omics data, preprocessing, models development, evaluation and biomarker discovery. A key aspect of playOmics is to make the results of predictive modeling as clear and understandable as possible, achieved through comprehensive visualizations and the application of SHAP values. PlayOmics is developed as an opensource and scalable tool, suitable for application in research as well as clinical environments. Its functionalities include the discovery of new genetic markers and conducting molecular diagnosis through the use of pre-trained models.

It adheres to the principles of open science, emphasizing reproducibility, repeatability and transparency, supported by its logging capabilities and an intuive graphical interface. Its dockerized setup simplifies the usage, aiming to make data analysis accessible.

By implementing a targeted pruning strategy that excludes poorly performing models early on, we significantly lowered the computational resources needed for our analysis.

Among other cutting-edge tools for multi-omics analysis, QLattice stands out as the most similar, both functionally and methodologically. Both tools offer valuable insights into biomedical data analysis, each with distinct strengths. QLattice impresses with its speed and capacity to manage extensive feature sets, attributed to its genetic algorithm. However, the commercial nature of QLattice might limit its accessibility due to its closed-code approach. In contrast, playOmics sets itself apart with its capacity to handle missing data and provide a user-friendly end-to-end data streaming pipeline, which notably enhances research reproducibility and repeatability.

Although playOmics did not outperform QLattice in benchmarking exercise, it demonstrated superior results in the “use case” experiment, where the utilization of an extensive dataset yielded better outcomes (validation MCC of 0.773) than those presented in benchmark comparisons. It’s worth noting that running QLattice on the same dataset led to the exclusion of all entries with missing data, which greatly reduced the dataset size. This significant reduction in data contributed to a notable decline in QLattice’s overall effectiveness (data not shown). This showcases playOmics’ unique ability to leverage all available information for deeper insights and ability to gracefully handle missing data, extracting as much signals as possible.

The discussion around playOmics and its comparison with other algorithms highlights the critical balance between model complexity and interpretability. While autoML, utilizing algorithms like stacked ensemble and deep learning models, showcased remarkable predictive accuracy with the highest validation MCC of 0.764, their complexity often masks the underlying decision-making processes. This complexity contrasts with simpler models like decision trees, which, while easier to interpret, offered lower performance metrics, evidenced by a validation MCC of 0.572. This contrast was further illuminated when comparing the performance of models built with a larger set of features versus those constructed from a smaller, more focused set. Models utilizing a more concise set of top 5 features across all data achieved higher validation MCC scores than models aggregating the top 10 features from each dataset (resulting in 40 features). This finding emphasizes that a limited set of features can enhance both the performance and interpretability of models, aligning with the strategies adopted in playOmics and QLattice. It’s important to note that the results for these two packages represent average values across the ten best-performing models, potentially leading to lower apparent performance compared to single-model assessments of autoML, Lasso and decision trees.

The benchmark results across different feature selection methods—AUC, MIM, and MRMR—show varied impacts on model performance. Across all algorithms, validation MCC scores ranged broadly from 0.512 to 0.771, with autoML models achieving top performance at 0.771 using MRMR. This variation highlights the important role of careful feature selection in optimizing model outcomes, suggesting that the choice of method can significantly influence overall model performance.

The playOmics methodology is designed to identify a range of highly effective models, with the primary objective of uncovering biomarkers and hypothesizing about potential relationships within the data. Rather than prescribing the “best” model for the user to select, playOmics encourages exploration and discovery. To date, we offer no definitive guidance on selecting the optimal model. While choosing the highest-performing model may seem straightforward, we allow to consider data availability in clinical environment. This is important, because due to various reasons, such as different detection capabilities or specific study limitations that preclude certain tests, not all omics data and variables might be available. Therefore, playOmics promotes a more nuanced and flexible approach to model selection, highlighting the importance of adaptability in research activities.

## Methods

### Metrics available in playOmics setup

playOmics provides a suite of metrics to comprehensively assess the predictive power of experiments.

The Matthews Correlation Coefficient (MCC) was selected as a primary metric for result evaluation in this paper. MCC has been described as a robust metric, especially in the context of imbalanced datasets [35]. MCC measures the quality of classification. A higher MCC value denotes superior performance, with the scale ranging from -1 (total disagreement) through 0 (no better than random chance) to +1 (perfect prediction). The MCC is calculated as follows:

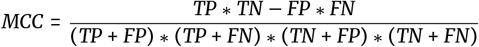

Additional metrics available include: accuracy, ROC AUC, Negative Predictive Value (NPV), Positive Predictive Value (PPV), precision, recall, sensitivity, specificity, and the F measure. These metrics, alongside counts of observations from each class predicted correctly or incorrectly for both validation and training data, equip users with detailed insights into model performance.

### Ranking methodology

To assess the performance of the playOmics framework relative to other analytical methods, a comparative study was conducted involving the QLattice package, autoML tool, Lasso regression, and decision tree models. The findings from this comparative analysis are presented in Table 2.

**Table 2.**
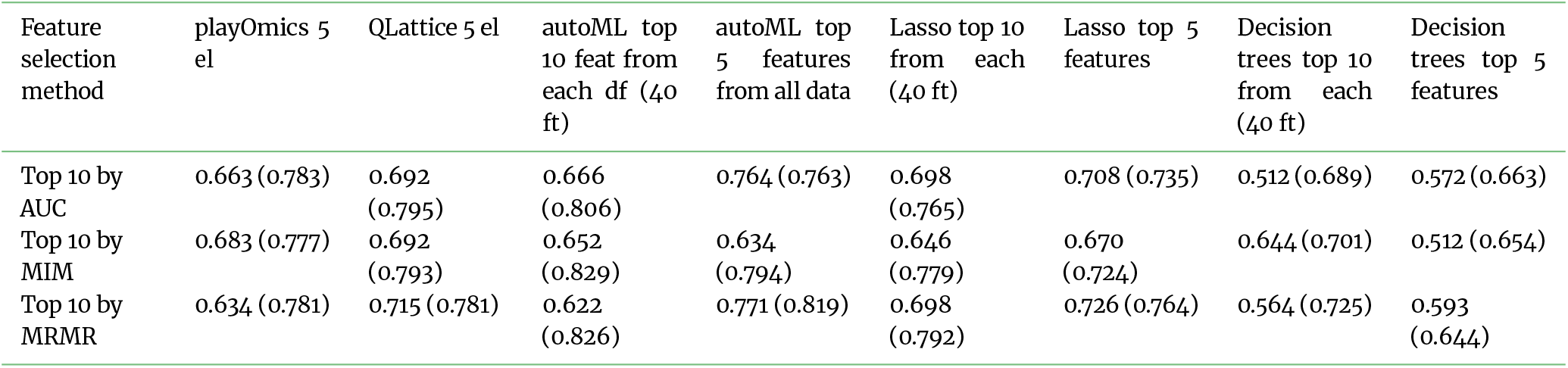
Performance comparison of playOmics versus other algorithms for BRCA subtypes (ductal/lobular) prediction on a limited dataset. Mean values of validation MCC metrics for 10 best models are presented for playOmics and QLattice. For other algorithms, the performance of the best model is shown. The values within brackets represent metrics obtained during training. AutoML, Lasso and decision trees were evaluated under two different scenarios: first, using top 10 features per dataset and second, using top 5 features across all datasets, to mimic the small models produced by playOmic and QLattice. Three different feature selection methods were applied, each selecting 10 features with highest class separation capacity. “n.a.” - did not return a result.

| Feature selection method | playOmics 5 el | QLattice 5 el | autoML top 10 feat from each df (40 ft) | autoML top 5 features from all data | Lasso top 10 from each (40 ft) | Lasso top 5 features | Decision trees top 10 from each (40 ft) | Decision trees top 5 features |
| --- | --- | --- | --- | --- | --- | --- | --- | --- |
| Top 10 by AUC | 0.663 (0.783) | 0.692 (0.795) | 0.666 (0.806) | 0.764 (0.763) | 0.698 (0.765) | 0.708 (0.735) | 0.512 (0.689) | 0.572 (0.663) |
| Top 10 by MIM | 0.683 (0.777) | 0.692 (0.793) | 0.652 (0.829) | 0.634 (0.794) | 0.646 (0.779) | 0.670 (0.724) | 0.644 (0.701) | 0.512 (0.654) |
| Top 10 by MRMR | 0.634 (0.781) | 0.715 (0.781) | 0.622 (0.826) | 0.771 (0.819) | 0.698 (0.792) | 0.726 (0.764) | 0.564 (0.725) | 0.593 (0.644) |

Our evaluation focused on distinguishing between BRCA subtypes (ductal versus lobular). Feature selection for the initial dataset for each target was executed, employing three distinct metrics in the univariate manner: Area Under the Curve (AUC), Mutual Information Maximization (MIM), and Minimum Redundancy Maximum Relevance (MRMR).

For the purpose of comparison, we adopted the Matthews Correlation Coefficient (MCC) metric. For both playOmics and QLattice, we assessed the performance of the top 10 models selected via training MCC metrics, subsequently calculating and presenting their mean value for the validation dataset, presented in table. For the remaining algorithms (autoML, Lasso, and Decision Tree), the analysis was conducted on the single best-performing model, identified by the highest MCC score on the training data.

Our methodology further specifies two distinct approaches for autoML, Lasso, and Decision Tree algorithms. Initially, each algorithm was fed with the top 10 best features identified from each dataset. Subsequently, to align with the comparative framework established for QLattice and playOmics, we refined our approach by selecting only the top 5 features across all datasets to input into these algorithms.

### Computation environment

All results presented in the article, except those for QLattice algorithm, were generated within a dockerized RStudio container hosted on a computing server equipped with 40 cores and 128 GB of RAM. The QLattice results were obtained from executing code provided by the Abzu team, which is accessible on GitHub, within a Python environment.

## Availability of supporting source code and requirements

Project name: playOmics

Project home page: https://github.com/JagGlo/playOmics

Operating system(s): Platform independent

Programming language: R

Other requirements: -

License: GNU GPL 3

## Data Availability

The data sets supporting the results of this article are available within a Docker image that can be downloaded from [36].

## Additional Files

### Abbreviations

AUC: Area Under the Curve
autoML: Automated Machine Learning
BRCA: Breast Cancer dataset from The Cancer Genome Atlas
CCA: Canonical Correlation Analysis
CNV: Copy Number Variation
F1: F1 Score
LDA: Linear Discriminant Analysis
MCC: Matthews Correlation Coefficient
MIM: Mutual Information Maximization
miRNA: MicroRNA
MRMR: Minimum Redundancy Maximum Relevance
NPV: Negative Predictive Value
PCA: Principal Component Analysis
PPV: Positive Predictive Value
RNA-Seq: RNA Sequencing
ROC: Receiver Operating Characteristic
SHAP: SHapley Additive exPlanations
TCGA: The Cancer Genome Atlas.

## Authors’ Contributions

J.G-W, K.S., K.W., T.G.: methodology design; J.G-W, K.S.: implementation; J.G-W: data analysis and manuscript preparation; T.G.-revision and supervision. All authors read and contributed to the manuscript.

## Acknowledgement

The playOmics package was created as part of the follow-up activities in EPISTOP project. Authors would like to thank the EPISTOP researchers and medical professionals for their cooperation and critical insights, which transformed the initial concept into fully equipped tool. Laura Bąkała suggested the change from MLflow to native “ object logging.

## Author notes

## Supplementary Data

